# Gut feelings: host identity shapes microbiomes in cartilaginous fishes

**DOI:** 10.64898/2026.11.24.766244

**Authors:** Thomas Hénault, Diana S. Vasconcelos, Teresa Moura, Catarina Maia, Raquel Xavier, Ana Veríssimo

## Abstract

This study provides the first characterization of the gut microbiome for several cartilaginous fishes and assesses the influence of host systematics and ecology on microbial diversity and composition. Cloacal swabs from 100 individuals representing 14 species of sharks, rays, and a chimaera were collected along the Portuguese coast across coastal, shelf and deep-water habitats (90–760 m depth). Gut bacterial communities were analysed using 16S rRNA gene metabarcoding targeting the V4 region. The results showed that microbiome diversity and structure varied significantly among host species, and that host identity is the main driver of gut microbiome composition. In contrast, habitat had no significant effect, although coastal species showed a tendency towards higher microbial diversity than deep-water taxa. Two phyla, Bacillota and Pseudomonadota were the dominant bacterial phyla across nearly all species, although their relative abundances differed between sharks vs. rays. In particular, three deep-water sharks displayed distinct and less diverse microbiomes dominated by a single bacterial genus, suggesting a stronger host–microbiome specialization. Functional predictions revealed differences in metabolic pathways associated with habitat and host systematics, particularly in nutrient processing and carbohydrate metabolism. Overall, these findings provide new insights into host–microbiome associations in this ancient vertebrate group, and highlight the importance of host-specific factors in shaping gut microbial communities.

## Introduction

Human-driven decline of many long-lived aquatic species, such as cartilaginous fishes (chimaeras, sharks and rays) is now well-documented and alarming (Dulvy et al. 2014; Pacoureau et al. 2021). To slow down their decline, there’s an urgent need to improve conservation strategies supported by reliable indicators of population health and ecological resilience (Kindsvater et al. 2018). In this context, microbiome studies could serve as valuable tools for monitoring population health and ecological resilience, reflecting changes driven by both biotic and abiotic factors (Sehnal et al. 2021).

Animal microbiomes are defined as the community of microorganisms (including prokaryotes, viruses and fungi), their genomes and the functional traits they encode (Marchesi et al. 2015; Berg et al. 2020). Over evolutionary time, hosts and their microbiome have developed symbiotic relationships, in which microorganisms provide essential metabolic and protective functions while benefiting from a stable habitat and nutrient supply (Chow et al. 2010). Consequently, the microbiome has emerged as a key factor in animal adaptation to diverse environments and offers valuable insights into an animal’s health status (Apprill et al. 2017). This growing recognition has led to a rapid expansion of microbiome studies in marine species, with derived data supporting the crucial role these microbial communities play in host health, ecology, and evolution (Brugman et al. 2018).

Across marine species, particularly in teleost fishes, the gut microbiome has been shown to vary with both biotic factors, such as host systematics, age, disease, and trophic level, and abiotic factors, including habitat type and environmental conditions (Wang et al. 2018; Egerton et al. 2018; Xavier et al. 2020; Kim et al. 2021; Rosado et al. 2023). Importantly, there is limited recruitment from bacterioplankton, indicating strong host-selective factors (Sylvain and Derome, 2017). Fish microbiome research has largely focused on teleost species, especially on commercial or aquaculture species (Xavier et al. 2024). In contrast, cartilaginous fishes have received comparatively little attention (but see Pogoreutz et al. 2019; Doane et al., 2020; Montemagno et al. 2024), despite their ecological importance in marine ecosystems (Ferretti et al. 2010; Barley et al. 2017; Shipley et al. 2023). Only recently, a few studies have begun to investigate the microbiomes of this group, and particularly of sharks, aiming to better understand how species identity, ecology, and behaviour shape their microbial communities. Like other marine vertebrates, the gut microbiomes of cartilaginous fishes appear to follow comparable ecological and host-related patterns (Johri et al. 2019; Perry et al. 2021). Host-specific effects have been reported, with bacterial communities differing even among individuals of the same species and also from bacterioplankton in the water (Storo et al. 2021, Bregman et al. 2023; Montemagno et al. 2024). Species-specific differences have also been documented in shark species, with some species displaying more distinct microbial communities than others, potentially due to ecological or evolutionary factors, such as host systematics (Storo et al. 2021; Black et al. 2021; Lynn & Wenzel, 2026). Environmental context can play a crucial role in shaping cartilaginous microbiome: comparisons between wild and captive ray populations have shown that wild individuals consistently exhibit higher bacterial diversity and distinct community composition, with pollution also having a negative impact on the gut microbiota of sharks eliciting microbial imbalance (Pinnell et al. 2021; Clavere-Graciette et al. 2022; Bregman et al. 2023). Although data on cartilaginous fish microbiome is growing, integrative comparative approaches across species with contrasting ecological strategies (e.g. habitat) and evolutionary histories are still largely lacking (Pratte et al. 2022).

In this study, we characterize for the first time the gut microbiomes of 14 cartilaginous species, including holocephalan and elasmobranch (sharks and rays) taxa. Our main aim was to provide a better understanding of how the gut microbiome varies across species and according to host systematics and ecology. For that, 100 individuals from 13 elasmobranch (8 sharks and 5 rays) and one chimaera species were sampled along the Portuguese coast from 90 to 760 m deep, covering coastal, shelf and deep-water habitats. Microbiome samples were collected using cloacal swabs and characterized using 16S rRNA (V4 region) metabarcoding. Specifically, we investigated whether microbiome composition and structure differ with host systematics and ecological traits.

## Materials and Methods

### Sample collection and processing

Cloaca swab samples from various cartilaginous fish species were collected in 2021 and 2022 during two annual trawl surveys coordinated by the Portuguese Institute for the Sea and Atmosphere (IPMA), under the National Biological Sampling Program (PNAB), as part of the EU/DGMARE Fisheries’ Data Collection Framework (DCF): the Nephrops Survey Offshore Portugal and the Portuguese International Bottom Trawl Survey (PT-IBTS). Both surveys were conducted onboard the R/V Mário Ruivo in June/July and October/November, respectively. A total of 100 individuals belonging to 14 species were sampled in the scope of this work, including sharks *Deania profundorum* (Smith & Radcliffe 1912)*, Etmopterus spinax* (Linnaeus 1758)*, Etmopterus pusillus* (Lowe 1839)*, Galeus atlanticus* (Vaillant 1888)*, Galeus melastomus* (Rafinesque 1810)*, Scyliorhinus canicula* (Linnaeus 1758)*, Scymnodon ringens* (Barbosa du Bocage & de Brito Capello 1864) and *Squalus blainville* (Risso 1827); rays *Dipturus oxyrinchus* (Linnaeus 1758)*, Leucoraja naevus* (Müller & Henle 1841)*, Raja circularis* (Couch 1838)*, Raja clavata* (Linnaeus 1758), and *Raja montagui* (Fowler 1910); and the chimaera *Chimaera monstrosa* (Linnaeus 1758) (Electronic supplementary material, Table S1). All individuals were brought onboard, quickly sampled for cloacal swabs, measured for total length and sexed. Swabs were placed in sterile Ziplock bags and stored at −20 °C upon collection. Cloacal swabs were used as a non-invasive proxy for the gut microbiome, an approach previously applied in elasmobranchs (Perry et al. 2021; Pinnell et al. 2021; Storo et al. 2021; Clavere-Graciette et al. 2022; Pratte et al. 2022; Bregman et al. 2023; Montemagno et al. 2024; Lyon et al. 2024). However, cloacal communities may also represent a transition between external and internal microbiomes (Perry et al. 2021). Metadata compiled for each species included systematics (species and phylogenetic lineage: sharks vs. rays vs. chimaeras) and ecological traits (habitat: coastal vs deep-water) (electronic supplementary material table S1). The habitat was attributed based on each species ecological affinity. Coastal species (*R. circularis, R clavata, R. montagui, S. blainville, S. canicula* and *L. naevus*) were considered those known to preferentially inhabit waters from shoreline to 200 m deep whereas deep-waters species (*C. monstrosa, D. profundorum, E, spinax, E. pusillus, G. atlanticus, G. melastomus, S. ringens* and *D. oxyrinchus*) were considered those that distribute preferentially deeper than 200 m.

### DNA extraction, library preparation, and sequencing

DNA from swabs was extracted using the DNeasy PowerSoil Pro Kit (Qiagen, Hilden, Germany), following the manufacturer’s instructions. One extraction blank was included for each batch of samples processed simultaneously (n = 6) and included in all downstream procedures. Extracted DNA was amplified by PCR targeting the V4 hypervariable region of the bacterial 16S rRNA gene (∼250 bp) using primers 515F (5′-GTGCCAGCMGCCGCGGTAA-3′) and 806R (5′-GGACTACHVGGGTWTCTAAT-3′) that target Archaea and Bacteria (Apprill et al. 2015; Parada et al. 2016). Each PCR reaction contained 5 µL of QIAGEN Multiplex PCR Master Mix (Qiagen; Crawley; UK), 0.3 µL of each primer, 1.5 µL of DNA template, and nuclease-free water to a final volume of 10 µL. Negative PCR controls and one mock community (ZymoBIOMICS Microbial Community DNA Standard, Zymo Research, USA) were processed along with the bacterial DNA samples and extraction blanks. PCRs were performed with the following program: 94 °C for 3 min; 35 cycles of 94 °C for 30 s, 50 °C for 1 min, and 72 °C for 1 min; followed by a final step at 72 °C for 10 min. Every sample was purified with AMPure XP beads at approximately 0.85× ratio to remove primer dimers (Beckman Coulter Genomics). A second PCR was performed using a distinct combination of index sequences per sample using 7 µL of 2x KAPA HiFi, 2.8 µL of nuclease-free water, 2.8 µL of PCR product and 1.4 µL of index primer (P7/P5), with the following program: 95 °C for 3 min; 9 cycles of 95 °C for 30 s, 55 °C for 30 s, and 72 °C for 30 s; followed by a final step at 72 °C for 5 min. Sample concentrations were estimated using an Epoch™ Microplate Spectrophotometer (BioTek Instruments, Inc.; Winooski, VT, USA) and pooled in equimolar amounts. After pooling, fragment length and concentration were assessed with TapeStation 4200 High Sensitivity D1000 Assay (Agilent, Santa Clara, CA). Final library validation and quantification were performed with KAPA Library Quant Kit qPCR Mix (Bio-Rad iCycler). The library was sequenced with an Illumina MiSeq desktop sequencer using the MiSeq Reagent Kit v.3, 2x 250bp aiming at 25 000 paired end reads per sample.

### Bioinformatic processing

All analyses were performed using R software v4.5.1 (R Core Team 2025). After demultiplexing, 16S rRNA amplicons were denoised using DADA2 pipeline (Callahan et al. 2016). Primers were removed using *trimLeft* based on primer length. Read quality filtering was then performed using the *filterAndTrim* function, with sequences truncated at 220 bp (forward) and 180 bp (reverse) based on inspection of quality profiles. Chimeras were detected and removed. The final ASV table was constructed, and taxonomy was assigned using SILVA 138 v2 database (Quast et al. 2012) with *assignTaxonomy* function in DADA2. Potential contaminants were removed using extraction and PCR blanks and the *decontam* v1.28.0 package with prevalence method (Davis et al. 2017), and mock communities were analysed and compared to the bacterial composition provided by the manufacturer to assess amplification and sequencing biases. Samples with fewer than 1,000 reads after denoising were removed (n = 6). All ASVs identified only at the kingdom level, or unclassified at the kingdom and phylum levels, were removed. ASV table is provided as electronic supplementary material table S2. Read counts were normalized with the DESeq2 median-of-ratios method (*estimateSizeFactors* function), using user-supplied per-ASV geometric means calculated over non-zero counts ensuring correction for differences in sequencing depth while handling sparse microbial data (Love et al. 2014).

### Statistical analysis

The *phyloseq* v1.52.0 package (McMurdie and Holmes, 2013) was used to estimate and visualize the abundances of the ten most abundant bacterial phyla and genera, represented by more than 1% of relative abundances across the entire dataset. Relative abundances were calculated using the *transform_sample_counts* function.

We used Shannon and Faith’s Phylogenetic Diversity (PD) indexes as estimates of alpha-diversity using *estimate_richness* and the *pd* functions in *phyloseq* and *picante* packages v1.8.2 (Kembel et al. 2010), respectively. These two metrics provide different perspectives on species-level diversity, namely Shannon diversity combines species richness and relative abundances, while Faith’s PD also incorporates phylogenetic diversity of communities. To test whether alpha diversity metrics differed among hosts, we performed linear models using host systematics: namely species and phylogenetic lineage (sharks vs. rays vs. chimaera); and habitat (coastal vs. deep waters) as predictors. Model selection was conducted based on the Akaike Information Criterion (AIC) using the *stepAIC* function from the *MASS* package v7.3-65 (Venables & Ripley, 2009). Model assumptions were checked with residual diagnostics based on simulated residuals with *DHARMa* package v0.5.0 (Hartig, 2026), which test for deviations from the expected distribution, dispersion and outliers. Faith’s PD was log-transformed to meet model assumptions, whereas Shannon diversity showed no major departure from them.

The significance of predictors was then assessed using the *anova* function. Pairwise differences among species were tested with Tukey’s HSD using *tukey_hsd* function, from *rstatix* package (Kassambara, 2026), and significant groupings are shown as compact letter displays in the figures. Beta-diversity was estimated using the *vegan* package v2.7-1 (Oksanen et al. 2025) and using Bray-Curtis dissimilarities calculated with the *vegdist* function. Principal Coordinates Analysis (PCoA) was used for visualization of dissimilarities between samples using the *cmdscale* function. A distance-based Redundancy Analysis (dbRDA) was performed to assess the influence of host species, phylogenetic lineage and habitat as predictors on bacterial community composition, using the *capscale* function. To identify which factors best explained differences in community assemblage, the *ordistep* function was applied using a forward selection with 999 permutations. Selected model significance was assessed using *anova* function with 999 permutations. Separate analyses were conducted for the full dataset, and for deepwater and coastal species separately. Homogeneity of multivariate dispersion among host species was assessed using PERMDISP using *betadisper* function, based on Bray–Curtis dissimilarities. Permutation tests were performed using 999 permutations. Given the small and unequal sample sizes among species, the sensitivity of dispersion estimates to small-sample bias was additionally assessed using the bias-adjusted procedure implemented in *betadisper*.

Only species represented by more than three individuals were included in statistical models to ensure sufficient statistical power. For this reason, *R. montagui*, *L. naevus*, *D. profundorum*, and *S. ringens* were excluded from these models due to low sample sizes.

### Analysis of predicted function

To understand whether host systematics or habitat potentially influence microbiome function, we used the software Phylogenetic Investigation of Communities by Reconstruction of Unobserved States software (*PICRUSt2*) (Bolyen et al. 2019; Douglas et al. 2020) to predict bacterial metabolic functions. The weighted nearest sequenced taxon index (NSTI), which measures the phylogenetic distance between observed taxa and sequenced reference genomes, was used using default cutoff values. Predicted functions were collapsed using the Kyoto Encyclopedia of Genes and Genomes (KEGG) pathway metadata (Kanehisa et al. 2019). Given the compositional differences observed between deepwater and coastal shark species, DESeq2 differential abundance test was used on predicted KEGG Orthology (KO) counts to detect differences in predicted metabolic function between sharks from deepwater and coastal habitats, and between phylogenetic lineage (rays vs sharks vs chimaeras). P-values were adjusted for multiple testing using the Benjamini–Hochberg false discovery rate (FDR) correction. Pathways with an adjusted p-value < 0.05 were considered significantly different between habitats. Principal component analysis (PCA) was used to visualise among-sample variation in predicted KEGG pathway profiles.

## Results

Illumina sequencing of all samples generated total of 7,158,167 reads. After bioinformatic processing (including decontamination steps), 4,817,380 reads corresponding to 6,768 ASVs were used for downstream analyses. A total of 2,476 ASVs were observed across deep-water species and 4,292 ASVs across coastal species.

### Gut microbiome diversity in cartilaginous fishes

Alpha diversity levels were highly variable among cartilaginous fish taxa within and across phylogenetic lineages (sharks, rays, and chimaeras) and habitats (coastal vs. deepwater) (Fig. 1). The only significant source of variation in alpha diversity values was host species, regardless of the metric used (linear models: Shannon, F_(9,76)_ = 10.38; p < 0.001; log-transformed Faith’s PD, F_(9,76)_ = 10.17; p < 0.001). Both alpha diversity metrics were lowest in the deepwater sharks *E. spinax* and *G. atlanticus*, which differed significantly from *R. clavata*, *S. canicula*, *L. naevus*, and the deepwater species *C. monstrosa* and *D. oxyrinchus* (Tukey’s HSD, p < 0.05; Fig. 1). Habitat and phylogenetic lineage were collinear with species and could not be tested independently of it. Nevertheless, coastal species tended to show higher alpha diversity than deepwater species (Fig. 1). Considering microbiome beta diversity across cartilaginous fish, most species showed generally similar microbiota community structure except for three deepwater species *E. spinax*, *E. pusilus,* and *G. atlanticus*. These species showed markedly distinct signals from each other as well as from the remaining taxa (Fig.2), and *E. spinax* and *G. atlanticus* also had the lowest alpha diversity values (Fig. 1). To further explore patterns within each habitat, data were analysed separately for deepwater vs. coastal species using dbRDA and visualized with PCoA (Supplementary Fig.1a). While no new pattern was apparent among the deepwater taxa, coastal taxa showed slight differences between sharks (*S. blainville* and *S. canicula*), whose values were spread over the PCoA1 axis, and the skate *R. clavata*, whose values were spread over the PCoA2 axis (Supplementary Fig. 1b). Host species was the only predictor retained by forward model selection and significantly explained variation in bacterial community composition among all species (dbRDA: F_(9,76)_ = 2.31, p ≤ 0.001), among deepwater species (dbRDA: F_(5,48)_ = 3.11, p ≤ 0.001) and among coastal species (dbRDA: F_(3,28)_ = 1.20, p = 0.002). Multivariate dispersion differed weakly among species (PERMDISP: F_(9,76)_ = 2.00, p = 0.048). Unadjusted pairwise comparisons were significant only for pairs involving *E. spinax*, and none remained significant after correction for multiple testing (Benjamini–Hochberg-adjusted p ≥ 0.32). Dispersion heterogeneity was no longer significant when *E. spinax* was excluded (PERMDISP: F_(8,66)_ = 1.88, p = 0.071), and the species effect on community composition persisted (dbRDA: F_(8,66)_ = 1.93, p ≤ 0.001). These results suggest that the species-associated differences in bacterial community composition are robust to the exclusion of *E. spinax*, although some contribution of interspecific differences in dispersion cannot be ruled out.

**Figure 1:**
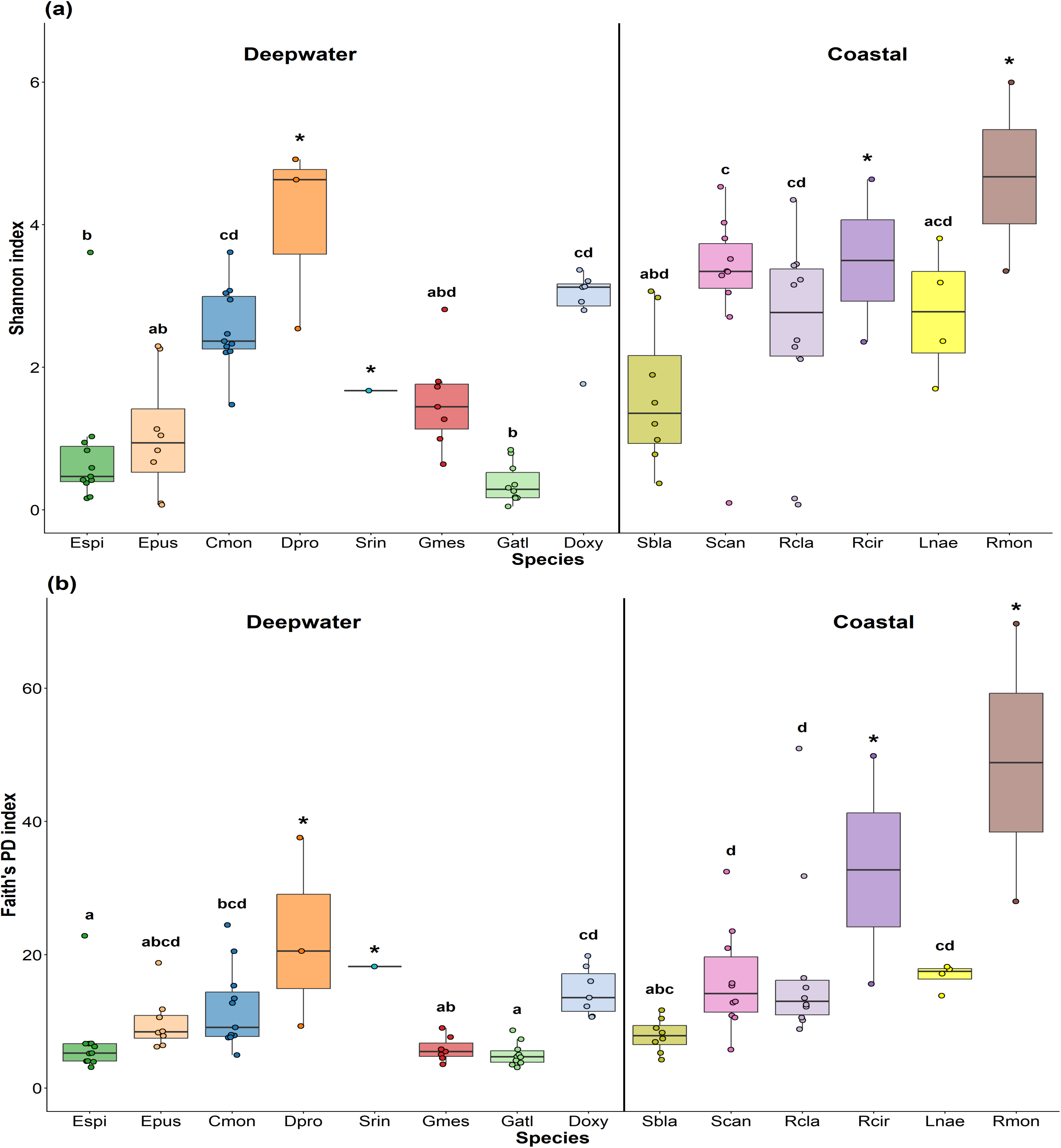
Boxplots of microbiota alpha diversity indices for deep-water and coastal species: (a) Shannon index and (b) Faith’s Phylogenetic Diversity (PD). Each colour represents a species. Different letters indicate significant differences among species according to Tukey’s HSD test (p < 0.05). Species sharing at least one letter are not significantly different. * Indicates species not included in pairwise comparisons due to low number of samples. Species codes are detailed in Supplementary Table S1.

**Figure 2:**
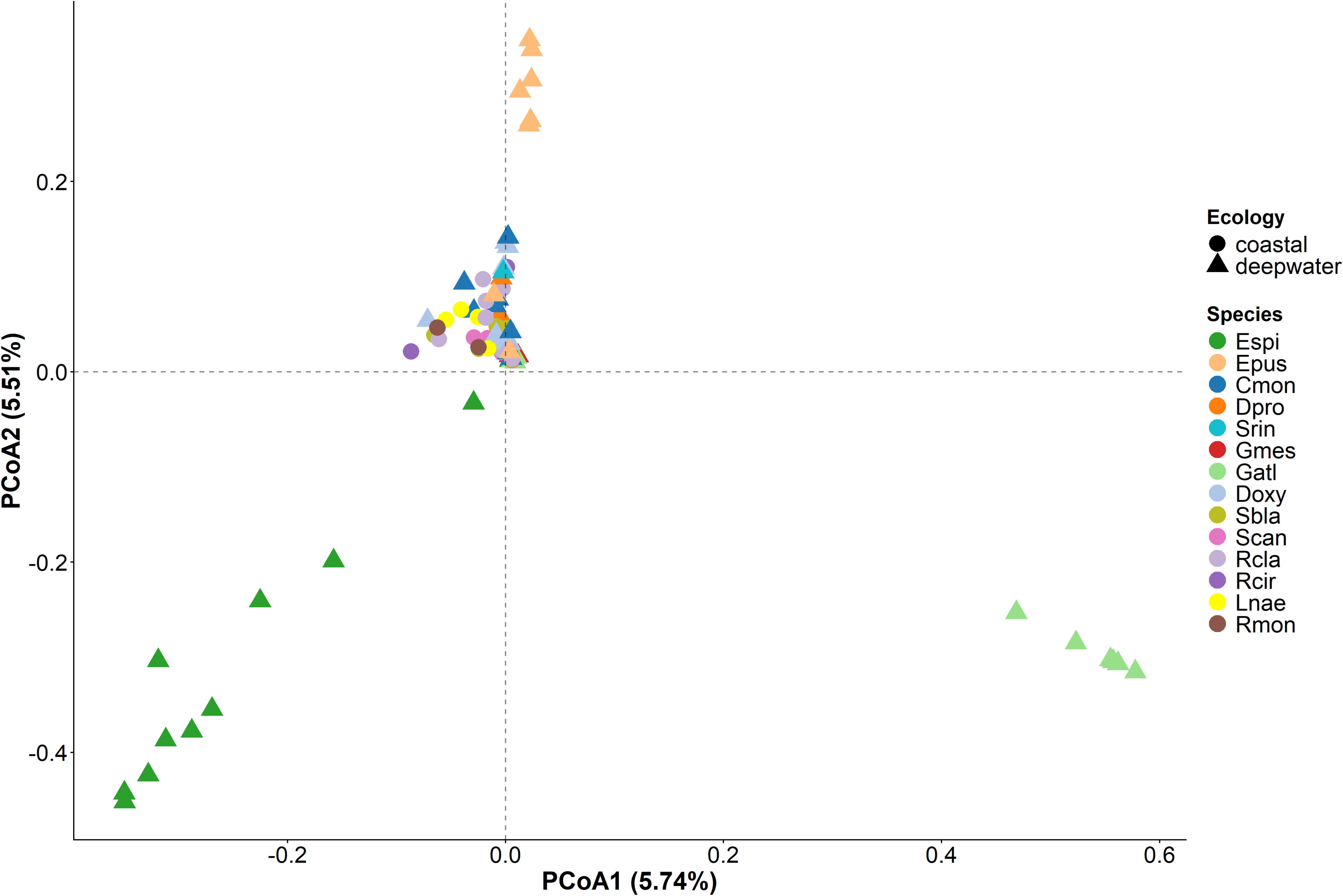
Principal Coordinates Analysis (PCoA) based on Bray–Curtis dissimilarities illustrating differences in bacterial community structure among individuals. Individual data points are coded by colour according to species, and by shape according to ecology (coastal or deepwater). Species codes are detailed in supplementary Table S1.

### Gut microbiome composition among cartilaginous fishes

The 10 most abundant phyla accounted for nearly 100% of the total microbiome composition in all species (Figure 3). In particular, two bacterial phyla, Bacillota and Pseudomonadota, consistently dominated the microbiomes of most cartilaginous fish species and collectively accounted for approximately 80–100% of the total gut microbial community depending on the host species. Bacillota was generally more abundant among sharks while the microbiome was dominated by Pseudomonadota in rays, regardless of habitat. Only *S. blainville* deviated from this pattern, with phylum Spirochaetota being predominant. However, results showed no significant differences in the structure of the most abundant phyla in rays and sharks. The only chimaera studied showed a phylum composition similar to rays although with increased contributions of Cyanobateriota and Planctomycetota.

**Figure 3:**
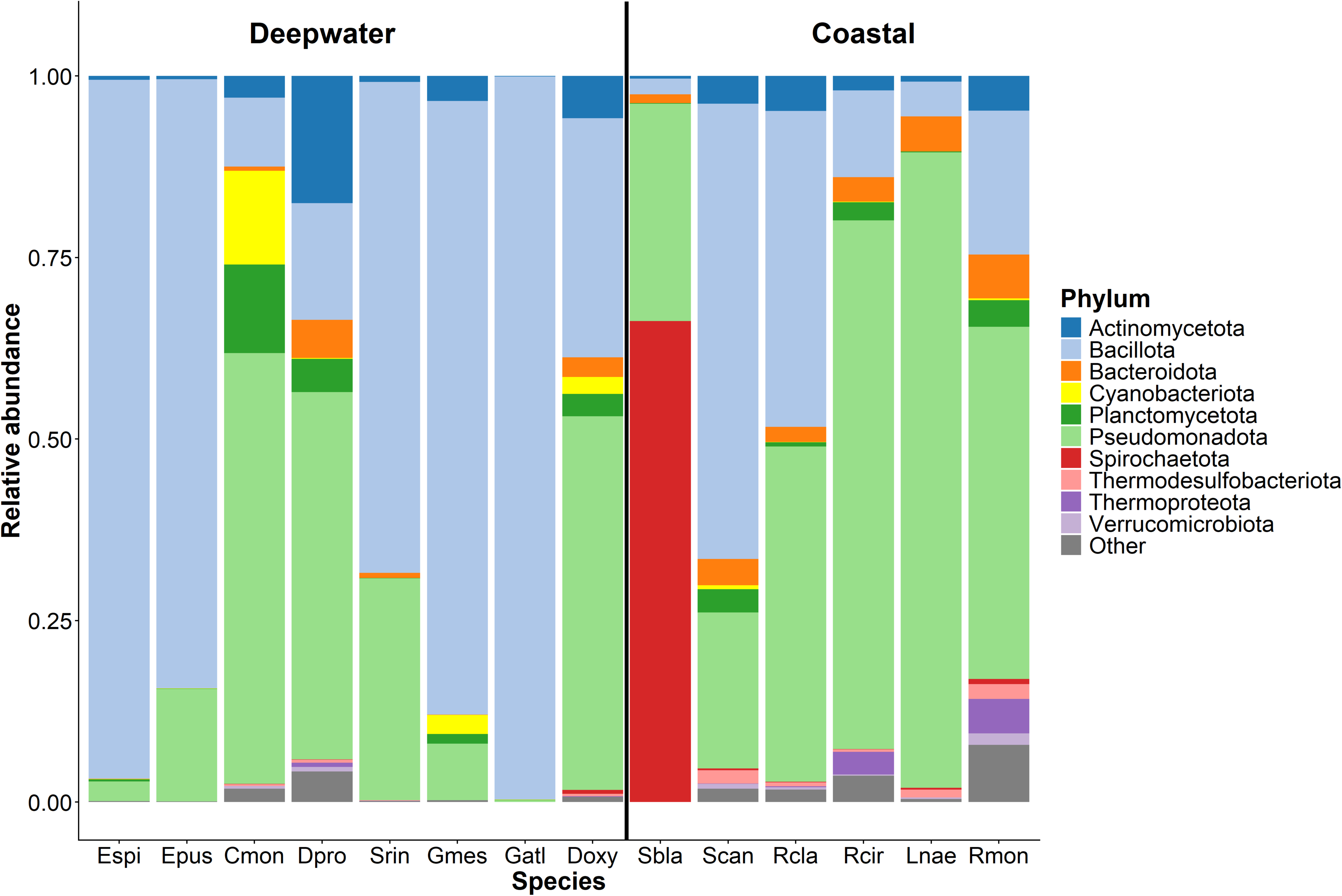
Stacked barplot showing relative abundance of the 10 most dominant bacterial phyla across species. Black line separates deep-water and coastal species. Species codes are detailed in Supplementary Table S1.

Genus-level analysis showed generally high inter-individual variability in bacterial genus composition and relative abundance (Figure 4), with this variability being generally higher in rays than in sharks (except in *S. canicula*). Notably, rays showed no clear dominance of a single bacterial genus, with each species and each individual host having distinct microbiome compositions. In turn, four deepwater shark species were strongly dominated by a single bacterial genus and showed low intra-specific variation. This was the case of *G. atlanticus* which was almost entirely dominated by *Mycoplasma,* and the case of both *Etmopterus* species and *G. melastomus* which were characterized by clear predominance of *Tyzzerella*. *Mycoplasma* was also present in small proportions in all *E. spinax* individuals, whereas *Psychrobacter* was detected among all *E. pusillus* individuals. *G. melastomus* had a small contribution of *Lactococcus*. On the other hand, the coastal *S. blainville* exhibited the predominance of two genera, *Brevinema* and *Photobacterium*, whose relative proportions varied among individuals. The chimaera *C. monstrosa* showed more consistent genus-level composition among individuals although the fraction of each bacterial genus varied considerably.

**Figure 4:**
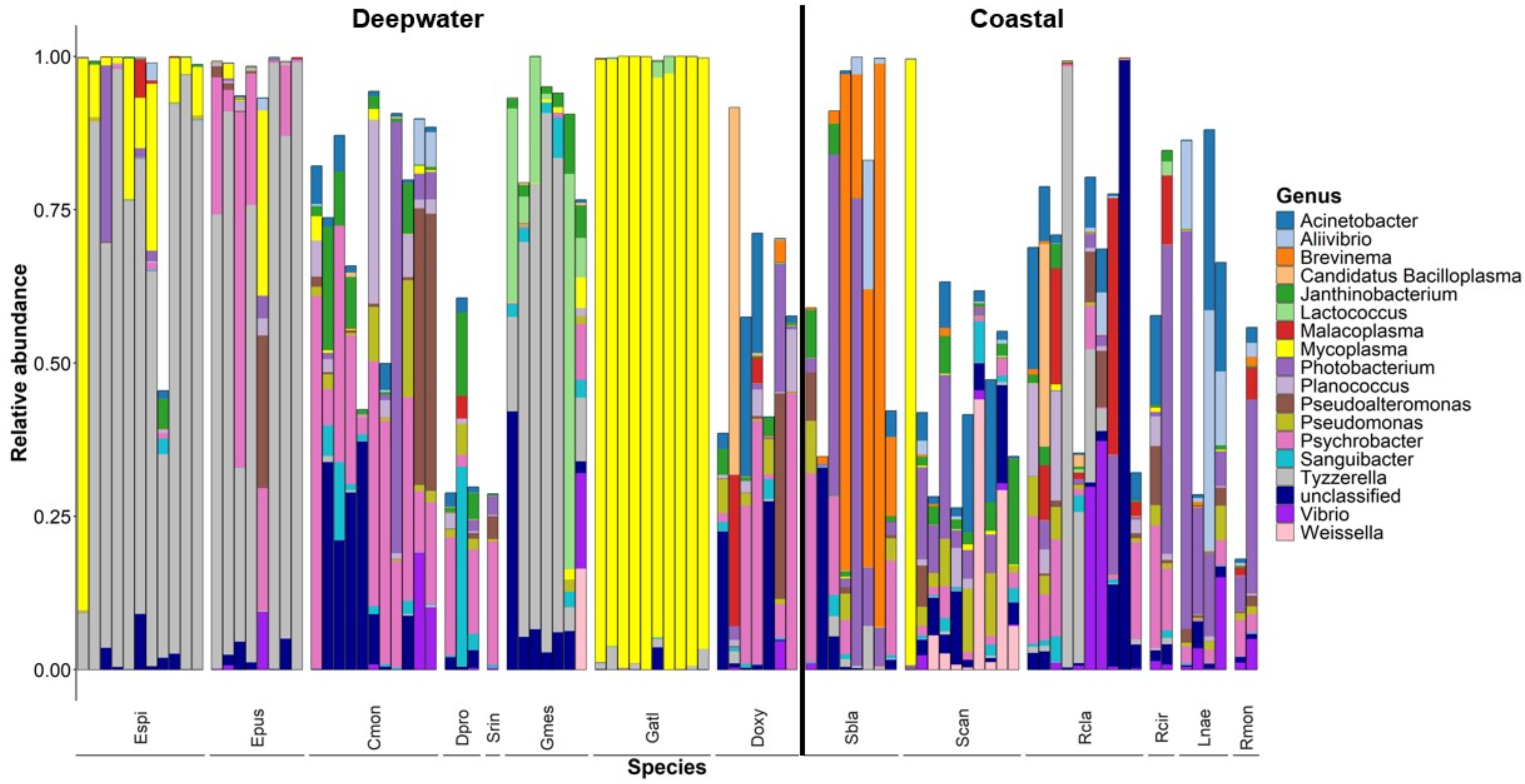
Stacked bar plot showing relative abundance of the most abundant bacterial genera (represented by > 1% of reads across all individuals) in the species included in the present study. Species codes are detailed in Supplementary Table S1.

### Functional profiling of cartilaginous fish gut microbiota

Analysis of predicted functional profiles revealed potential differences between coastal and deepwater sharks and between phylogenetic lineages. A total of 82 pathways were differentially abundant between coastal and deepwater species (64 were successfully annotated), whereas 72 pathways were differentially abundant between sharks and rays (53 were successfully annotated) (electronic supplementary material, tables S3 and S4), with the vast majority (only two exceptions) being enriched in sharks relatively to rays. Still, the majority of the predicted metabolic pathways enriched in the microbiome of sharks were also enriched in deepwater species. These corresponded to about 68% of the enriched pathways found in shark microbiome, and to 58% of the enriched pathways found in the microbiome of deepwater species, indicating association between the two factors that could be linked to unbalanced taxa sampling in the different habitats. Differences in the predicted microbial functions between deep and coastal water species and between rays and sharks were mostly related to nutrient processing and absorption (digestive system), and to metabolism of co-factors, vitamins and carbohydrate metabolism (electronic supplementary material, tables S3 and S4).

## Discussion

This study aimed to characterize for the first time the gut microbiota of different cartilaginous fish and investigate whether microbiome diversity varies according to host taxonomy (host species), phylogenetic lineage (shark vs rays vs chimaeras) and ecology (habitat). We show that host taxonomy modulates gut microbiome diversity, while host phylogeny and habitat have considerably smaller effects. Across marine species, and especially fish, gut microbiome diversity has been shown to vary according to host-related factors, including species identity and ecological niche (Huang et al. 2020). The marked variation in gut microbiota composition among cartilaginous fish species could reflect several interacting factors. Species differences in gut anatomy, mucus composition and immune function may act as host-filtering mechanisms and could be determinant for microbial colonisation and persistence (Zhao et al. 2026). In addition, fish species occupying different habitats may be exposed to distinct environmental microbial communities and conditions that influence microbiome assembly (Singh et al. 2025). On the other hand, differences in prey preference likely affect the nutrient availability in the gut. However, host diet may covary with host species, as well as with other factors such as host age and habitat occupancy, making their individual effects difficult to disentangle.

Our results showed that coastal species, albeit not significantly, exhibited higher intraindividual (alpha) microbial diversity, including highest phylogenetic heterogeneity in gut bacteria, compared to deep-water species. This could be a reflection of broader ecological niches, linked to a higher heterogeneity of habitats and preys in coastal environments. Additionally, the gut microbiome of coastal sharks included in this study exhibited more similarities between them, than to the coastal rays. Similarly, a previous study by Lyons et al. (2024) showed that the gut microbiomes of rays were more similar to each other than to that of sharks. This is indicative that phylogenetic position within cartilaginous fish, and in particular differences between rays and sharks, could also be contributing towards gut microbiome diversity in the group. Our results suggest that the broader ecological distinction between deepwater and coastal species is insufficient to explain the observed microbial variability, and that other traits, which may or may not be entirely linked to habitat preferences, play a role in shaping gut microbial assemblages of cartilaginous fish. Interestingly, our ordination analyses of interindividual dissimilarities revealed that three deepwater sharks (*E. spinax*, *E. pusilus* and *G. atlanticus*) exhibited distinct gut microbiome structures when compared with the remaining species, with high levels of divergence found between them as well. Notably, these three species also exhibited lower microbial diversity which could arise from specialization to the lower temperatures, high hydrostatic pressure and/or lower nutrient availability in deep-sea environments (Webb et al. 2010; Osman and Weinnig, 2022). A potential lower capacity to recruit a diverse gut microbial community or increased specialization, was hypothesized to lead to stronger host–microbiome associations in deepwater species under extreme and resource-limited conditions (Osman and Weinnig, 2022). Previous observations have indeed reported that organisms gut microbiome diversity tends to decrease with increasing depth, becoming more specialized and reflecting the constrained and selective conditions of deep-sea environments (Ohwada et al. 1980; Osman and Weinnig, 2022). Feeding constraints in deep-water environments, including reduced prey availability and lower feeding rates, may also contribute to the lower gut microbiome diversity observed in these deep-water species (Draven et al. 2017). Low diversity and symbiotic microbiome in deep sea teleost fish has been documented in anglerfish (Freed et al. 2019). Importantly, the differentiation of the deepwater sharks *Etmopterus* spp and *G. atlanticus*, seems to stem from a strong dominance by *Tyzerella* spp. Interestingly, the dominance of *Mycoplasma* in *G. atlanticus* is akin to the dominance of *Tyzzerella* in three other deep-water sharks, including in its congener. The genus *Mycoplasma* has not previously been reported as a dominant component of the gut microbiome of cartilaginous fishes, although it has been detected at low relative abundances in the bonnethead shark and white shark (Leigh et al. 2021; Pratte et al. 2022), neither of which are considered deep-water taxa. In turn, *Tyzzerella* has been reported from the gut of several marine organisms, including crustaceans (Shui et al. 2020), fish (Escalas et al. 2021) and elasmobranchs (Pratte et al. 2022; Clavere-Graciette et al. 2022). While *E. spinax*, *E. pusillus*, and *G. melastomus* are known to have a limited dietary breadth, share similar prey spectra, and may also occupy overlapping habitats (Rodríguez-Garcías et al. 2024; Nuño Muñoz, 2015), little information is available for *G. atlanticus*. It can be hypothesized that specific features of *G. atlanticus* may explain its unique dominance by *Mycoplasma.* Unfortunately, data on deepwater cartilaginous fishes’ ecology is still very limited, limiting our ability to infer which other important factors may be contributing to the gut microbiome heterogeneity found between deepwater species.

Examination of the taxonomic composition of the gut microbiome showed that two phyla dominated the gut microbiome of all cartilaginous fishes surveyed here: Bacillota and Pseudomonadota. These two phyla are present in high proportions in the gut microbiota of most vertebrates (Ley et al. 2008) and are also consistently reported as dominant components of the gut microbiome across a wide range of marine species, including teleost fish (Egerton et al. 2018). Accordingly, these two phyla have also been widely reported as major components of the gut microbiome of cartilaginous fishes, with Pseudomonadota frequently described as the dominant phylum in this group. Interestingly, shark species in our dataset exhibited a higher relative abundance of Bacillota compared to Pseudomonadota, whereas the opposite pattern was observed in rays and in the chimaera. Although Pseudomonadota is often reported as the dominant phylum in elasmobranch gut microbiomes, most previous studies focused on coastal species and match our observations in coastal rays (Clavere-Graciette et al. 2022; Montemagno et al. 2024). Our results further showed that the gut microbiome of deepwater sharks, can instead be dominated by phylum Bacillota. In other deepwater fish species, Pseudomonadota has been reported as the dominant phylum in gut microbiomes (Collins et al. 2021; Le Cam Ligier et al. 2026). These contrasting patterns suggest that differences in gut microbiome composition may not be explained by habitat depth but may also reflect host-specific characteristics and ecological factors. In contrast, individuals of *S. blainville* were mostly dominated by a different phylum, Spirochaetota, mainly represented by the genus *Brevinema*. This pattern was not consistent across individuals, some of which were instead dominated by other genera, such as *Photobacterium*. To our knowledge, the dominance of Spirochaetota has not previously been reported in cartilaginous fishes. *Brevinema* has been detected in the gut microbiota of several teleosts, including salmonids (Brown et al. 2019; Steiner et al. 2022), gilthead sea bream (Domingo-Bretón et al. 2025) and European sea bass (Alfonso et al. 2023), and has been described as a potential opportunistic taxon in rainbow trout (Brown et al. 2019). Its abundance has been reported to increase with rising water temperature in farmed fish (Steiner et al. 2022; Domingo-Bretón et al. 2025). In gilthead sea bream, this shift was not driven by colonisation from the surrounding water and was associated with an imbalance in intestinal homeostasis (Domingo-Bretón et al. 2025). The interindividual variability in *Brevinema* dominance suggests that it may be influenced by individual-specific factors, such as diet, physiological state or environmental exposure, although these potential drivers could not be investigated in our study.

As discussed above, analysis of microbiome composition at the genus level also showed differences between deep-water sharks and the other chondrichthyans. The gut microbiomes of deep-water sharks exhibited lower variability, particularly in *G. atlanticus* and *Etmopterus* spp., consistent with the strong dominance of a single bacterial genus. Analysis of predicted functional profiles further suggested ecological differentiation between gut bacteria of cartilaginous fish from coastal and deep-water environments, particularly in pathways related to digestive processes and vitamin metabolism.

Predicted functional profiles also suggested differences between rays and sharks. However, these patterns may be strongly influenced by the greater representation of deep-water sharks in the dataset, as most metabolic pathways enriched in sharks were also enriched in deep-water species. However, these results need to be interpreted with caution since predictions of microbial functional potential were inferred from 16S rRNA gene sequences based on the phylogenetic relationships between detected bacteria and reference genomes. The limited representation of fish-associated bacterial lineages in reference databases is likely to limit the accuracy of these predictions. Moreover, predicted pathway enrichment does not necessarily reflect differences in gene expression or actual metabolic activity. Nevertheless, these results allow the posing of new hypothesis regarding symbiotic nature of host-microbiome associations, that may reflect different adaptations between tested groups. Although our results showed no significant differences in microbiome composition diversity between rays and sharks, and between deepwater and coastal species, future studies should aim to disentangle the influence of these variables using a more balanced sampling design of phylogenetic lineage and habitat.

In conclusion, we found that host species is the main driver of the gut microbiome diversity in cartilaginous fishes. Many ecological and functional hypotheses, such as the role of diet and habitat in shaping the gut microbiome of cartilaginous fish, warrant further investigation. While our findings provide new insights into host–microbiome interactions in cartilaginous fish, they should be interpreted with caution given the small sample sizes, narrow phylogenetic and ecological breadth. Future studies should aim for improved representation of the remarkable cartilaginous fish host diversity, and further in-depth testing of the possible effect of life stage, depth and sampling location on the microbiome.

## Acknowledgments

The authors are grateful to the scientific team onboard the RV Mário Ruivo, for their assistance in sample collection.

## Author Contributions

AV and RX contributed to the study conception and design. Sampling and material preparation were performed by TM and CM; laboratory work was performed by TH and DSV; data collection and analysis were performed by all authors. The first draft of the manuscript was written by TH, AV and RX, and all authors commented on previous versions of the manuscript. All authors read and approved the final manuscript.

## Ethical Statement

Ethical approval was not required for the studies involving animals in accordance with the local legislation and institutional requirements because animals were sacrificed as part of routine biological sampling programmes run by Instituto Português do Mar e Atmosfera (IPMA) All procedures were performed by a certified individual and following EU laws, as described in the Ethics statements and remaining methods section.

## Data availability statement

All data used in this manuscript is provided as supplementary materials. Raw sequences will be made publicly available upon acceptance in NCBI’s Sequence Read Archive.

## Conflict of interest

The authors declare no conflicts of interest

## Funding

Samples were collected in the scope of the Portuguese Biological Sampling Program under the EU data Collection Framework (PNAB/DCF) (MAR-03.02.01-FEAMP-0016). TH was provided by the program “*Service Civique”* administered by Mission Locale Caen Le Mer Calvados Centre. This work was supported by FCT - Fundação para a Ciência e Tecnologia, I.P. by projects with reference 2022.02937.CEECIND/CP1730/CT0011 (to AV), and 2020.00854.CEECIND/CP1601/CT0001 (to RX). The authors have no relevant financial or non-financial interests to disclose.

## Notes

### Competing Interest Statement

The authors have declared no competing interest.

## References

Alfonso, S., Mente, E., Fiocchi, E. et al (2023) Growth performance, gut microbiota composition, health and welfare of European sea bass (*Dicentrarchus labrax*) fed an environmentally and economically sustainable low marine protein diet in sea cages. Sci Rep. 10.1038/s41598-023-48533-3

Apprill A (2017) Marine Animal Microbiomes: Toward Understanding Host–Microbiome Interactions in a Changing Ocean. Front Mar Sci. 10.3389/fmars.2017.00222

Apprill A, McNally S, Parsons R, Weber L (2015) Minor revision to V4 region SSU rRNA 806R gene primer greatly increases detection of S*AR11* bacterioplankton. Aquat Microb Ecol. 10.3354/ame01753

Barley S, Meekan M, Meeuwig J (2017) Species diversity, abundance, biomass, size and trophic structure of fish on coral reefs in relation to shark abundance. Mar Ecol Prog Ser. 10.3354/meps11981

Berg G, Rybakova D, Fischer D et al (2020) Microbiome definition re-visited: Old concepts and new challenges. Microbiome. 10.1186/s40168-020-00875-0

Bolyen E, Rideout JR, Dillon MR et al (2019) Reproducible, interactive, scalable and extensible microbiome data science using QIIME 2. Nat Biotechnol. 10.1038/s41587-019-0209-9

Bregman G, Lalzar M, Livne L, Bigal E, Zemah-Shamir Z, Morick D, Tchernov D, Scheinin A and Meron D (2023) Preliminary study of shark microbiota at a unique mix-species shark aggregation site, in the Eastern Mediterranean Sea. Frot Microbiol. 10.3389/fmicb.2023.1027804

Brugman S, Ikeda-Ohtsubo W, Braber S, Folkerts G, Pieterse CMJ, Bakker PAHM (2018) A Comparative Review on Microbiota Manipulation: Lessons From Fish, Plants, Livestock, and Human Research. Front Nutr. 10.3389/fnut.2018.00080

Callahan BJ, McMurdie PJ, Rosen MJ, Han AW, Johnson AJA, Holmes SP (2016) DADA2: High-resolution sample inference from Illumina amplicon data. Nat Methods. 10.1038/nmeth.3869

Black C, Merly L, Hammerschlag N (2021) Bacterial Communities in Multiple Tissues Across the Body Surface of Three Coastal Shark Species. Zool Stud. 10.6620/ZS.2021.60-69

Brown, R. M., Wiens, G. D., & Salinas, I. (2019) Analysis of the gut and gill microbiome of resistant and susceptible lines of rainbow trout (Oncorhynchus mykiss). Fish & shellfish immunology. 10.1016/j.fsi.2018.11.079

Chiarello M, Villéger S, Bouvier C, Auguet JC, Bouvier T (2017) Captive bottlenose dolphins and killer whales harbor a species-specific skin microbiota that varies among individuals. Sci Rep. 10.1038/s41598-017-15220-z

Chow J, Tang H, Mazmanian SK (2011) Pathobionts of the gastrointestinal microbiota and inflammatory disease. Curr Opin Immunol. 10.1016/j.coi.2011.07.010

Clavere-Graciette AG, McWhirt ME, Hoopes LA, Bassos-Hull K, Wilkinson KA, Stewart FJ, Pratte ZA (2022) Microbiome differences between wild and aquarium whitespotted eagle rays (*Aetobatus narinari*). Anim Microbiome. 10.1186/s42523-022-00187-8

Collins FWJ, Walsh CJ, Gomez-Sala B et al (2021) The microbiome of deep-sea fish reveals new microbial species and a sparsity of antibiotic resistance genes. Gut Microbes. 10.1080/19490976.2021.1921924

Davis NM, Proctor DM, Holmes SP, Relman DA, Callahan BJ (2017) Simple statistical identification and removal of contaminant sequences in marker-gene and metagenomics data. Microbiome. 10.1101/221499

Degregori S, Schiettekatte NMD, Casey JM, Brandl SJ, Mercière A, Amato KR, Mazel F, Parravicini V, Barber PH (2024) Host diet drives gut microbiome convergence between coral reef fishes and mammals. Mol Ecol. 10.1111/mec.17520

Delport TC, Power ML, Harcourt RG, Webster KN, Tetu SG (2016) Colony Location and Captivity Influence the Gut Microbial Community Composition of the Australian Sea Lion (*Neophoca cinerea*). Appl Environ Microbiol. 10.1128/AEM.00192-16

Doane MP, Morris MM, Papudeshi B et al (2020) The skin microbiome of elasmobranchs follows phylosymbiosis, but in teleost fishes, the microbiomes converge. Microbiome. 10.1186/s40168-020-00840-x

Domingo-Bretón R, Cools S, Moroni F, Belenguer A, Calduch-Giner JA, Croes E, Holhorea PG, Naya-Català F, Boon H, Pérez-Sánchez J (2025) Intestinal microbiota shifts by dietary intervention during extreme heat summer episodes in farmed gilthead sea bream (*Sparus aurata*). Aquac Rep. 10.1016/j.aqrep.2024.102566

Douglas GM, Maffei VJ, Zaneveld JR, Yurgel SN, Brown JR, Taylor CM, Huttenhower C, Langille MGI (2020) PICRUSt2 for prediction of metagenome functions. Nat Biotechnol. 10.1038/s41587-020-0548-6

Drazen JC, Sutton TT (2017) Dining in the Deep: The Feeding Ecology of Deep-Sea Fishes. Ann Rev Mar Sci. 10.1146/annurev-marine-010816-060543

Dulvy NK, Fowler SL, Musick JA et al (2014) Extinction risk and conservation of the world’s sharks and rays. eLife. 10.7554/eLife.00590

Egerton S, Culloty S, Whooley J, Stanton C, Ross RP (2018) The Gut Microbiota of Marine Fish. Front Microbiol. 10.3389/fmicb.2018.00873

Erwin PM, Rhodes RG, Kiser KB, Keenan-Bateman TF, McLellan WA, Pabst DA (2017) High diversity and unique composition of gut microbiomes in pygmy (*Kogia breviceps*) and dwarf (*K. sima*) sperm whales. Sci Rep. 10.1038/s41598-017-07425-z

Escalas A, Auguet J-C, Avouac A, Seguin R, Gradel A, Borrossi L, Villéger S (2021) Ecological Specialization Within a Carnivorous Fish Family Is Supported by a Herbivorous Microbiome Shaped by a Combination of Gut Traits and Specific Diet. Front Mar Sci. 10.3389/fmars.2021.622883

Fadiel A (2007) Mycoplasma genomics: Tailoring the genome for minimal life requirements through reductive evolution. Front Biosci. 10.2741/2207

Farias I, Figueiredo I, Moura T, Serrano Gordo L, Neves A, Serra-Pereira B (2006) Diet comparison of four ray species (*Raja clavata, Raja brachyura*, Raja montagui and Leucoraja naevus) caught along the Portuguese continental shelf. Aquat Living Res. 10.1051/alr:2006010

Ferretti F, Worm B, Britten GL, Heithaus MR, Lotze HK (2010) Patterns and ecosystem consequences of shark declines in the ocean. Ecol Lett. 10.1111/j.1461-0248.2010.01489.x

Freed, L. L., Easson, C., Baker, L. J., Fenolio, D., Sutton, T. T., Khan, Y., Blackwelder, P., Hendry, T. A., & Lopez, J. V. (2019). Characterization of the microbiome and bioluminescent symbionts across life stages of ceratioid anglerfishes of the Gulf of Mexico. FEMS Microbiology Ecology. 10.1093/femsec/fiz146

Hartig F (2026). DHARMa: Residual Diagnostics for Hierarchical (Multi-Level / Mixed) Regression Models. R package version 0.5.0, https://github.com/florianhartig/dharma.

Johri S, Doane M, Allen L, Dinsdale E (2019) Taking Advantage of the Genomics Revolution for Monitoring and Conservation of Chondrichthyan Populations. Diversity. 10.3390/d11040049

Kanehisa M, Sato Y, Furumichi M, Morishima K, Tanabe M (2019) New approach for understanding genome variations in KEGG. Nucleic Acids Res. 10.1093/nar/gky962

Kassambara A (2026) *rstatix: Pipe-Friendly Framework for Basic Statistical Tests*. R package version 1.1.0, https://rpkgs.datanovia.com/rstatix/.

Kembel SW, Cowan PD, Helmus MR, Cornwell WK, Morlon H, Ackerly DD, Blomberg SP, Webb CO (2010) Picante: R tools for integrating phylogenies and ecology. Bioinformatics. 10.1093/bioinformatics/btq166

Kim PS, Shin N-R, Lee J-B, Kim M-S, Whon TW, Hyun D-W, Yun J-H, Jung M-J, Kim JY, Bae J-W (2021) Host habitat is the major determinant of the gut microbiome of fish. Microbiome. 10.1186/s40168-021-01113-x

Kindsvater HK, Dulvy NK, Horswill C, Juan-Jordá M-J, Mangel M, Matthiopoulos J (2018) Overcoming the Data Crisis in Biodiversity Conservation. Trends Ecol Evol. 10.1016/j.tree.2018.06.004

Kousteni V, Karachle PK, Megalofonou P (2017) Diet and trophic level of the longnose spurdog *Squalus blainville* (Risso, 1826) in the deep waters of the Aegean Sea. Deep-Sea Res I: Oceanogr Res Pap. 10.1016/j.dsr.2017.04.008

Le Cam Ligier, C., Garel, M., Thibault, H., Guasco, S., Tamburini, C., Ménard, F., Cherel, Y., Casalot, L., & Martini, S. (2026) Bacterial composition of the microbiome of mesopelagic fishes from the Northeast Atlantic. BMC biology. 10.1186/s12915-026-02677-6

Leigh SC, Papastamatiou YP, German DP (2021) Gut microbial diversity and digestive function of an omnivorous shark. Mar Biol. 10.1007/s00227-021-03866-3

Ley RE, Hamady M, Lozupone C, Turnbaugh PJ, Ramey RR, Bircher JS, Schlegel ML, Tucker TA, Schrenzel MD, Knight R, Gordon JI (2008) Evolution of Mammals and Their Gut Microbes. Science. 10.1126/science.1155725

Llewellyn MS, McGinnity P, Dionne M, Letourneau J, Thonier F, Carvalho GR, Creer S, Derome N (2016) The biogeography of the atlantic salmon ( *Salmo salar* ) gut microbiome. ISME J. 10.1038/ismej.2015.189

Love MI, Huber W, Anders S (2014) Moderated estimation of fold change and dispersion for RNA-seq data with DESeq2. Genome Biol. 10.1186/s13059-014-0550-8

Lynn ID, Wenzel MA (2026) Ephemeral signatures of phylosymbiosis in dermal microbiomes within the requiem shark family (Carcharhinidae). Access Microbiol. 10.1099/acmi.0.001140.v2

Lyons K, Bedore CN, Carlisle AB, Moniz L, Odom TL, Ahmed R, Greiman SE, Freedman RM (2024) Network Analysis Reveals Species-Specific Organization of Microbial Communities in Four Co-Occurring Elasmobranch Species along the Georgia Coast. Fishes. 10.3390/fishes9010034

Marchesi JR, Ravel J (2015) The vocabulary of microbiome research: A proposal. Microbiome. 10.1186/s40168-015-0094-5

McMurdie PJ, Holmes S (2013) phyloseq: An R Package for Reproducible Interactive Analysis and Graphics of Microbiome Census Data. PLoS ONE. 10.1371/journal.pone.0061217

Montemagno F, Romano C, Bastoni D, Cordone A, De Castro O, Stefanni S, Sperone E, Giovannelli D (2024) Shark Microbiome Analysis Demonstrates Unique Microbial Communities in Two Distinct Mediterranean Sea Shark Species. Microorganisms. 10.3390/microorganisms12030557

Muñoz LN (2015) Feeding Ecology of Small Deep-Water Lanternsharks (*Etmopterus spinax* and *Etmopterus pusillus*) off the Algarve Coast. Dissertations & Theses. Universidade do Algarve, Portugal. https://www.proquest.com/openview/6b7aaca085c0dea26d73d72e018676db/1?pq-origsite=gscholar&cbl=2026366&diss=y

Ohwada K, Tabor PS, Colwell RR (1980) Species Composition and Barotolerance of Gut Microflora of Deep-Sea Benthic Macrofauna Collected at Various Depths in the Atlantic Ocean. Appl Environ Microbiol. 10.1128/aem.40.4.746-755.1980

Oksanen, J., Simpson, G. L., Blanchet, F. G., Kindt, R., Legendre, P., Minchin, P. R., O’Hara, R. B., Sólymos, P., Stevens, M. H. H., Szoecs, E., Wagner, H., &, et al. (2025) vegan: Community Ecology Package. R package version 2.7–1. https://CRAN.R-project.org/package=vegan

Osman EO, Weinnig AM (2022) Microbiomes and Obligate Symbiosis of Deep-Sea Animals. Ann Review Anim Biosci. 10.1146/annurev-animal-081621-112021

Pacoureau N, Rigby CL, Kyne PM et al (2021) Half a century of global decline in oceanic sharks and rays. Nature. 10.1038/s41586-020-03173-9

Parada AE, Needham DM, Fuhrman JA (2016) Every base matters: Assessing small subunit rRNA primers for marine microbiomes with mock communities, time series and global field samples. Environ Microbiol. 10.1111/1462-2920.13023

Perry CT, Pratte ZA, Clavere-Graciette A et al (2021) Elasmobranch microbiomes: Emerging patterns and implications for host health and ecology. Anim Microbiome. 10.1186/s42523-021-00121-4

Pinnell LJ, Oliaro FJ, Van Bonn W (2021) Host-associated microbiota of yellow stingrays (*Urobatis jamaicensis*) is shaped by their environment and life history. Mar Freshw Res. 10.1071/MF20107

Pogoreutz, C., Gore, M. A., Perna, G., Millar, C., Nestler, R., Ormond, R. F., Clarke, C. R., & Voolstra, C. R. (2019) Similar bacterial communities on healthy and injured skin of black tip reef sharks. Anim Microbiome. 10.1186/s42523-019-0011-5

Pratte ZA, Perry C, Dove ADM, Hoopes LA, Ritchie KB, Hueter RE, Fischer C, Newton AL, Stewart FJ (2022) Microbiome structure in large pelagic sharks with distinct feeding ecologies. Anim Microbiome. 10.1186/s42523-022-00168-x

Quast C, Pruesse E, Yilmaz P, Gerken J, Schweer T, Yarza P, Peplies J, Glöckner FO (2012) The SILVA ribosomal RNA gene database project: Improved data processing and web-based tools. Nucleic Acids Res. 10.1093/nar/gks1219

R Core Team (2025) R: A Language and Environment for Statistical Computing. R Foundation for Statistical Computing. https://www.R-project.org/

Rasmussen JA, Kiilerich P, Madhun AS, Waagbø R, Lock E-JR, Madsen L, Gilbert MTP, Kristiansen K, Limborg MT (2023) Co-diversification of an intestinal *Mycoplasma* and its salmonid host. ISME J. 10.1038/s41396-023-01379-z

Razin S, Yogev D, Naot Y (1998) Molecular Biology and Pathogenicity of Mycoplasmas. Microbiol. Mol. Biol. Rev. 10.1128/MMBR.62.4.1094-1156.1998

Ripley B, Venables B (2009) MASS: Support Functions and Datasets for Venables and Ripley’s MASS (p. 7.3–65) [Dataset]. 10.32614/CRAN.package.MASS

Rocha EPC (2002) Genomic repeats, genome plasticity and the dynamics of *Mycoplasma* evolution. Nucleic Acids Res. 10.1093/nar/30.9.2031

Rodríguez-García C, Gonçalves Neto JB, García-Romero C, Domínguez-Bustos ÁR, Cabrera-Castro R (2024) Feeding habits of two shark species: Velvet belly, Etmopterus spinax (Linnaeus, 1758) and blackmouth catshark, *Galeus melastomus* (Rafinesque, 1810), present in fishing discards in the Gulf of Cádiz. Environ Biol Fish. 10.1007/s10641-024-01519-0

Rosado D, Canada P, Marques Silva S, Ribeiro N, Diniz P, Xavier R (2023) Disruption of the skin, gill, and gut mucosae microbiome of gilthead seabream fingerlings after bacterial infection and antibiotic treatment. FEMS Microbes. 10.1093/femsmc/xtad011

Sehnal L, Brammer-Robbins E, Wormington AM, Blaha L, Bisesi J, Larkin I, Martyniuk CJ, Simonin M, Adamovsky O (2021) Microbiome Composition and Function in Aquatic Vertebrates: Small Organisms Making Big Impacts on Aquatic Animal Health. Front Microbiol. 10.3389/fmicb.2021.567408

Shipley ON, Matich P, Hussey NE et al (2023) Energetic connectivity of diverse elasmobranch populations – implications for ecological resilience. Proc R Soc B: Biol Sci. 10.1098/rspb.2023.0262

Shui Y, Guan Z-B, Liu G-F, Fan L-M (2020) Gut microbiota of red swamp crayfish Procambarus clarkii in integrated crayfish-rice cultivation model. AMB Express. 10.1186/s13568-019-0944-9

Singh, B. K., Thakur, K., Kumari, H., Mahajan, D., Sharma, D., Sharma, A. K., Kumar, S., Singh, B., Pankaj, P. P., & Kumar, R. (2025) A review on comparative analysis of marine and freshwater fish gut microbiomes: insights into environmental impact on gut microbiota. FEMS microbiology ecology. 10.1093/femsec/fiae169

Steiner K, Laroche O, Walker SP, Symonds JE (2022) Effects of water temperature on the gut microbiome and physiology of Chinook salmon (Oncorhynchus tshawytscha) reared in a freshwater recirculating system. Aquaculture. 10.1016/j.aquaculture.2022.738529

Storo R, Easson C, Shivji M, Lopez JV (2021) Microbiome Analyses Demonstrate Specific Communities Within Five Shark Species. Front Microbiol. 10.3389/fmicb.2021.605285

Sylvain, F. É., & Derome, N. (2017). Vertically and horizontally transmitted microbial symbionts shape the gut microbiota ontogenesis of a skin-mucus feeding discus fish progeny. Sci Rep. 10.1038/s41598-017-05662-w

Wang AR, Ran C, Ringø E, Zhou ZG (2018) Progress in fish gastrointestinal microbiota research. Reviews in Aquac. 10.1111/raq.12191

Webb TJ, Vanden Berghe E, O’Dor R (2010) Biodiversity’s Big Wet Secret: The Global Distribution of Marine Biological Records Reveals Chronic Under-Exploration of the Deep Pelagic Ocean. PLoS ONE. 10.1371/journal.pone.0010223

Xavier R, Pereira A, Pagan A, Hendrick GC, Nicholson MD, Rosado D, Soares MC, Pérez-Losada M, Sikkel PC (2020) The effects of environment and ontogeny on the skin microbiome of two *Stegastes* damselfishes (Pomacentridae) from the eastern Caribbean Sea. Mar Biol. 10.1007/s00227-020-03717-7

Xavier R, Severino R, Silva SM (2024) Signatures of dysbiosis in fish microbiomes in the context of aquaculture. Rev Aquac. 10.1111/raq.12862

Zhao, Y., Ren, Z., Xu, Q., Zhu, T., Hu, H., Fu, Y., Jiang, J., & Zhai, Q. (2026) Factors influencing gut microbial colonization: A host-microbe-environment interaction perspective. Current research in food science. 10.1016/j.crfs.2026.101361

